# From root to shoot; Quantifying nematode tolerance in *Arabidopsis thaliana* by high-throughput phenotyping of plant development

**DOI:** 10.1101/2023.03.15.532731

**Authors:** Jaap-Jan Willig, Devon Sonneveld, Joris J.M. van Steenbrugge, Laurens Deurhof, Casper C. van Schaik, Misghina G. Teklu, Aska Goverse, Jose L. Lozano-Torres, Geert Smant, Mark G. Sterken

## Abstract

Nematode migration, feeding site formation, withdrawal of plant assimilates, and activation of plant defence responses have a significant impact on plant growth and development. Plants display intraspecific variation in tolerance limits for root-feeding nematodes. Although disease tolerance has been recognised as a distinct trait in biotic interactions of mainly crops, we lack mechanistic insights. Progress is hampered by difficulties in quantification and laborious screening methods. We turned to the model plant *Arabidopsis thaliana*, since it offers extensive resources to study the molecular and cellular mechanisms underlying nematode-plant interactions. Through imaging of tolerance-related parameters the green canopy area was identified as an accessible and robust measure for assessing damage due to cyst nematode infection. Subsequently, a high-throughput phenotyping platform simultaneously measuring the green canopy area growth of 960 *A. thaliana* plants was developed. This platform can accurately measure cyst- and root-knot nematode tolerance limits in *A. thaliana* through classical modelling of tolerance limits. Furthermore, real-time monitoring provided data for a novel view of tolerance, identifying a compensatory growth response. These findings show that our phenotyping platform will enable further studies into a mechanistic understanding of tolerance to below-ground biotic stress.

**Highlight:** The mechanisms of tolerance to root-parasitic nematodes remain unknown. We developed a high-throughput phenotyping system that enables unravelling the underlying mechanisms of tolerance to nematodes.

## Introduction

The impact of root-parasitic nematodes on plant growth and development leads to reduced yield of food crops globally (Jones *et al*., 2013; Bebber *et al*., 2014). Interestingly, plants show intraspecific variation in their growth responses to belowground biotic stresses by nematodes (Miltner et al., 1991; Potter and Dale, 1994). In other words, plants display different disease burdens in response to similar initial nematode densities. The first measurable repression in plant growth (i.e., yield, plant height) by pathogen infection is defined as the tolerance limit (Seinhorst, 1985). The tolerance limit of plants is measured independently from resistance because tolerance and resistance are genetically and physiologically independent traits (Evans & Haydock, 1990; Teklu *et al*., 2022). The tolerance limit is dependent on the plant and the nematode species, and the interaction between those two. Even though nematode tolerance is known as a phenomenon for multiple decades, we lack an understanding of the underlying molecular and/or genetic mechanisms. Difficulties in quantifying and the required laborious screening methods hamper progress in this research area. Therefore, there is a need for a genetically tractable model species and a scalable screening method to better understand tolerance.

The model plant species should be able to support infections of the families of cyst nematodes (*Heteroderinae*) and root-knot nematodes (*Meloidogyninae*), as these belong to the most damaging obligate sedentary root parasites worldwide (Jones *et al*., 2013; Quist *et al*., 2015). Both genera spend most of their life cycle within roots. Cyst nematodes and root-knot nematodes cause significant damage to the root tissue by mechanical damage, large morphological changes, and loss of plant assimilates (Kyndt *et al*., 2013). Infective juveniles (J2s) of cyst and root-knot nematodes invade the plant root and move through different tissue layers to reach the vascular cylinder (Wyss & Zunke, 1986; Wyss *et al*., 1992). Cyst nematodes move brutally through cells causing significant mechanical damage, whereas root-knot nematodes move in between cells by gently pushing them aside. After reaching the vascular cylinder, cyst and root-knot nematodes induce a redifferentiation process that leads to the formation of multinucleated feeding cells known as syncytium and giant cells, respectively. The feeding cells are hypermetabolic sink tissues and serve as the sole source of nutrients for growing nematodes throughout their entire life cycle. This leads to less nutrients available for plant growth and development. Both nematode families consist of multiple species, of which the beet cyst nematode *Heterodera schachtii* and the root-knot nematode *Meloidogyne incognita* are the best studied, and therefore considered to be model species for plant-nematode interaction studies.

Up to now, tolerance to root-parasitic nematodes is mainly studied in field crops (Seinhorst, 1985; Norshie *et al*., 2011; Sasanelli *et al*., 2013; Been *et al*., 2015; Moosavi, 2015), making it difficult to resolve the underlying mechanisms as to why plants differ in their tolerance limits. Knowledge of the genetic underpinnings of tolerance is important for understanding how plants differ in their tolerance limits to root-feeding nematodes. The model plant *Arabidopsis thaliana* is a rich and often-used resource to study the role of natural genetic variation, molecular, and cellular mechanisms underlying nematode-plant interactions (Absmanner *et al*., 2013; Lozano-Torres *et al*., 2014; Anwer *et al*., 2018; Warmerdam *et al*., 2018). It has been shown that *A. thaliana* harbours significant natural genetic variation in susceptibility to the beet cyst nematode *Heterodera schachtii* and *Meloidogyne incognita in vitro* (Anwer *et al*., 2018; Warmerdam *et al*., 2018). For instance, a quantitative trait loci allele of *ATS40-3* that was identified with a Genome Wide Association Study affects the sex ratio of *H. schachtii* in Arabidopsis (Anwer *et al*., 2018). Recently, we also showed that the transcription factor TCP9 modulates tolerance to *H. schachtii* via reactive oxygen species mediated processes (Willig *et al*., 2022). However, it remains unknown if *A. thaliana* harbours significant genetic variation in tolerance to nematodes.

So far, the Seinhorst Yield Loss Model (SYLM) is mostly used to determine the tolerance limit for plant-parasitic nematodes (Seinhorst, 1985; Norshie *et al*., 2011; Sasanelli *et al*., 2013; Been *et al*., 2015; Moosavi, 2015). The SYLM models the density-yield relationship by an inverted sigmoid curve where loss of yield at a given timepoint is a derivative function of damage. Yield remains stable up to a certain nematode density i.e., the tolerance limit, then declines exponentially, to end at a minimum yield level at the highest nematode density. The SYLM enables a comparison of tolerance limits of different crop varieties for a specific pathogen (Seinhorst, 1985). Importantly, herein, yield loss is dependent on the number of parasitic nematodes in the soil and not the number of successful infections detected at harvest (Seinhorst, 1985). The tolerance limit and the steepness of the decline in yield over increasing nematode densities are both agronomically important traits. Alternatively, for other pathogens, the slope of a regression of the selected host parameter against the pathogen load are made to determine if a plant is more tolerant than the other (Pagán & Garcia-Arenal, 2020). However, both linear regression and the SYLM do not account for biological responses to pathogen infection like overcompensation or compensation (i.e., increased growth speed or growth speed recovery) (Pedigo *et al*., 1986).

Most of the studies that measure the effect of nematode infection on crop yield rely on destructive end-point measurements (*i*.*e*., crop yield (Sasanelli *et al*., 2013; Been *et al*., 2015)), leaving an information gap between start and endpoint of the experiment. Few time-series experiments were performed by measuring the effect of soybean cyst nematode *Heterodera glycines* on the root system by using a rhizotron (Miltner *et al*., 1991) or by measuring the effect of *Meloidogyne chitwoodi* densities on the height of potato plants (Norshie *et al*., 2011). Additionally, time-series measurements were also performed on the green canopy area upon nematode infection using a phenotyping design based on microplots (Joalland *et al*., 2016, 2017). However, this later design was limited to 70 microplots, which does accommodate large phenotyping experiments included hundreds of genotypes for multi-genotype comparisons (i.e., Genome-Wide Association Studies or mutant screens). A system that is able to monitor a larger number of individually measured plants over time could give insight in genes that modulate tolerance to root-parasitic nematodes.

Another returning hurdle in tolerance research is to determine which plant trait should be measured. Most of the studies on tolerance to root-parasitic nematodes in potato focus on yield reduction, e.g., potato yield expressed in tuber weight and quality (Been et al., 2015; Teklu et al., 2022), which depend on single destructive endpoint measurement. Others also measure the effect of nematodes foliage or plant height which are not perse destructive (Norshie *et al*., 2011), but these types of measurements are less often done because they are laborious. In *A. thaliana*, the green canopy growth, inflorescences, and the number of viable seeds can be used to measure tolerance to viruses (Shukla *et al*., 2018). Most of the screening methods are laborious and/or time-consuming. For many purposes, these kinds of measurements might be adequate, but for large scale phenotyping of multiple plant genotypes these laborious measurements hamper progress. Therefore, the chosen trait should also be easy to measure. However, it remains unknown which plant trait gives the most robust data for a high throughput design to quantify tolerance to root-parasitic nematodes in *A. thaliana*.

Here, we report on the construction and validation of a high-throughput phenotyping platform to monitor in real-time plant growth under belowground biotic stress caused by nematode infection. Initially, we tested growth parameters (e.g., root system architectural components, green canopy area, flowering, and silique formation) to robustly quantify the effect of *H. schachtii* infestation on the growth and development of the host. We found that the green canopy area of Arabidopsis is an accessible and informative trait for automated analysis of growth-responses to root-parasitic nematodes. Time-series measurements showed that daily growth rates of individual plants are congruent over plants and provided insight in biological responses of the host (*i*.*e*., compensation responses) that could not be determined in previous experimental designs. Therefore, we believe that our phenotyping system will not only allow for a more accurate mechanistic studies into tolerance, but can also ultimately pinpoint genes that modulate tolerance to biotic stress by endoparasitic nematodes.

## Materials and Methods

### Plant culturing

For seed sterilization, Arabidopsis seeds (Col-0, N60,000) were placed in Eppendorf tubes in a desiccator. The seeds were vapour sterilized for 3-4 hours using a mixture of hydrochloric acid (25%) and sodium hypochlorite (50 g/L). Finally, the sterile seeds were stratified for four days. For *in vitro* assays, seeds were sown on square petri dishes (120×120 mm) containing modified Knop medium (Sijmons *et al*., 1991). For *in vivo* pot experiments, non-sterilized seeds were also stratified for four days and sowed directly on top of silversand. Seedlings were grown at 21 °C and 16-h-light/8-h-dark conditions. Seedlings growing in our high-throughput platform were grown at 19 °C and 16-h-light/8-h-dark conditions with LED light (150 lumen).

### Hatching and sterilization of Heterodera schachtii and Meloidogyne incognita

*H. schachtii* cysts (Woensdrecht population from IRS, the Netherlands) were separated from sand and roots of infected *Brassica oleracea* plants as previously described (Baum et al., 2000). *H. schachtii* cysts were transferred into a clean Erlenmeyer containing water with 0.02% sodium azide. This mixture was gently stirred for 20 min. Later, sodium azide was removed by vigorously washing with tap water. Cysts were then placed on a hatching sieve in a glass petri dish. An antibiotic solution was added containing 1.5 mg/mL gentamycin, 0.05 mg/mL nystatin, and 3mM zinc chloride. The cysts were incubated in the dark for 4-7 days. Thereafter, nematode juveniles were collected in a 2 mL Eppendorf tube. For *in vitro* experiments, J2s were surface sterilized using a HgCl_2_-containing solution (0.002% Triton X-100 w/v, 0.004% NaN_3_ w/v, 0.004% HgCl_2_ w/v) for 20 min. After incubation, the nematodes were collected by centrifugation and the supernatant was removed. Nematodes were washed with sterile water and spun down again. This was repeated three times. Finally, for *in vitro* experiments, the nematodes were resuspended in 0.7% gelrite (Duchefa Biochemie, Haarlem, the Netherland). For *in vivo* experiments, hatched J2s were separated from debris by centrifugation in a 70% sucrose gradient. Afterwards, the nematodes were washed by centrifugation in tap water three times. Finally, the nematodes were resuspended in tap water.

Eggs of *Meloidogyne incognita* were obtained by soaking *M. incognita* (strain ‘Morelos’ from INRA, Sophia Antipolis, France) with 0.05% (v/v) NaOCl for three minutes. Roots were rinsed with tap water and the eggs were collected on a 25 µM sieve. Next, the eggs were incubated in a solution of 2.4 mM NaN_3_ for 20 minutes while swirling. Thereafter, the eggs were rinsed with tap water and incubated on a 25 µM sieve in a solution of 1.5 mg/mL gentamycin, 0.05 mg/mL nystatin in the dark at room temperature. After four days, hatched juveniles were collected. For *in vivo* experiments, hatched J2s were separated from debris by centrifugation in a 70% sucrose gradient. Afterwards, the nematodes were washed by centrifugation in tap water three times. Finally, the nematodes were resuspended in tap water.

### Quantifying the root system architecture of nematode-infected Arabidopsis

Nine-day-old *in vitro-grown Arabidopsis thaliana* seedlings were individually inoculated with increasing densities (*P*_*i*_)of surface sterilised second-stage juveniles of *H. schachtii* (from 0 to 50 juveniles per mL modified KNOP media). Roots of nematode-infected plants were scanned at 7 dpi using an Epson Perfection V800 photo scanner. Various root measurements were conducted (*i*.*e*. total root length, main root length, total secondary root length, and average secondary root length), collectively referred to as the root system architecture. Measurements were taken using the WinRHIZO package for Arabidopsis (WinRHIZO pro2015, Regent Instrument Inc., Quebec, Canada). The number of root tips was counted manually based on the scans made by WinRHIZO. Differences in the root length per seedling in centimetres and the number of root tips were statistically analysed with a one-way ANOVA with *post-hoc* Tukey’s HSD test in R.

### Quantifying above-ground development and growth of *Heterodera schachtii* infected Arabidopsis plants

Three-week-old *Arabidopsis thaliana* seedlings were inoculated with increasing densities of *H. schachtii* (from 0 to 50 juveniles per g dry sand). During a period of 14 days post inoculation (dpi), we recorded the green canopy area, the number of flowers, the number of siliques, and the number of basal stems every second day with a video camera. By using Adobe Photoshop (Version: 22.5.6 20220204.r.749 810e0a0 x64), we manually isolated the green canopy area from the images and analysed the area manually using ImageJ. Differences per treatment per timepoint were analysed using One-Way ANOVA with *post-hoc* Tukey’s HSD test in R.

### High throughput analysis of the green canopy area of nematode-infected Arabidopsis

Pots filled with silver sand were placed in stainless steel frames made (Fig. S1a and d) and covered with a 3 mm thick black nonreflective foamed PVC coversheet (Fig. S1b, c and e) drilled with countersunk holes ∼73 mm apart. Extra holes were drilled with a diameter of 7 mm and 4 mm away from the countersunk holes for nematode inoculations. Prior to sowing, silver sand was watered with Hyponex (1.7 mM/L NH4^+^, 4.1 mM/L K^+^, 2 mM/L Ca^2+^, 1.2 mM/L Mg^2+^, 4.3 mM/L NO_3-_, 3.3 mM/L SO_42-_, 1.3 mM/L H2PO_4-_, 3.4 µm/L Mn, 4.7 µm/L Zn, B 14 µm/L, 6.9 µm/L Cu, 0.5 µm/L Mo, 21 µm/L Fe, pH 5.8) for five minutes. Seven days after sowing, seedlings were watered again for five minutes. Nine-day-old seedlings were inoculated with increasing densities of *H. schachtii* and *M. incognita* (0 to 100 juveniles per g dry sand). Seedlings were inoculated by first punching a hole of approximately 3.5 cm deep in the soil, into which 1 mL tap water containing a measured number of nematodes was added using a pipet. Pictures of the Arabidopsis seedlings were taken every hour (15 pictures per day) during the light period automatically for a period of 21 days with twelve UI-1490LE-C-HQ cameras (IDS Imaging) mounted with 12mm lenses on the ceiling of the climate chamber (Cat. No. B5M12056, IDS Imaging). Because of the purple LED light, colour corrections were done using Adobe Photoshop (Version: 22.5.6 20220204.r.749 810e0a0 x64). The surface area of the rosette was determined using a custom-written ImageJ macro (ImageJ 1.51f; Java 1.8.0_321 [32-bit]) (Supplementary File 1) and Java was used to make GIFs (Supplementary File 2). More information about how the experiment was performed can be found in: (DOI: dx.doi.org/10.17504/protocols.io.kqdg39167g25/v1) (Private link for reviewers: https://www.protocols.io/private/C82F27F2BE7511ED91B00A58A9FEAC02 to be removed before publication.)

### Seinhorst yield loss model

To quantify the tolerance limit for *H. schachtii* infection in Arabidopsis we fitted the green canopy area data to the Seinhorst’s Equation (1) (Seinhorst, 1985). Before fitting the model to the data, each measurement was averaged over the day and replicates for each nematode density.

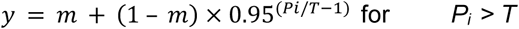

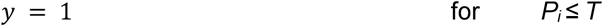

The parameters tolerance limits (*T*) and relative minimum yield (*m*) were estimated for all measurements.

### Plant growth analysis using the high-throughput phenotyping platform

To analyse the growth data of the plants obtained from the high-throughput platform, custom scripts and functions were written in “R” (available via gitlab: https://git.wur.nl/published_papers/willig_2023_camera-setup). For analysis we used the median daily leaf area (cm_2_), which was calculated by taking the median leaf area of the daily measurements (15 per day). The data was log_2_-transformed before analysis for normalization. The rate of growth was determined per day per plant by

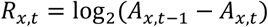

where *R* is the growth rate of plant *x* from day *t-1* to day *t* based on the median leaf surface area *A*. Differences in growth rate were determined using a Wilcoxon Rank Sum test as implemented in the ggpubr package (https://cran.r-project.org/web/packages/ggpubr/index.html).

Differences in growth rate between plants infected with either *H. schachtii* or *M. incognita* were tested using a paired t-test comparing data from the same day and density combination. Data was multiple-testing corrected using the *p*.*adjust* function with the *fdr* method (Benjamini & Hochberg, 1995).

### Modelling growth rates

Growth models were fitted to the data using the *growthrates* package, which was used to fit a logistic growth model to the data,

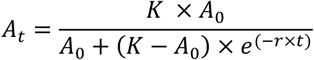

where the parameters *K* (maximum canopy area), *A*_*0*_ (initial canopy area, and *r* (intrinsic growth rate) were determined as a function of time *t* in days. These fitted values were used to explore the relation between nematode density and the *K* and *r*. Bad fits (p < 0.05) were removed before analysis.

We found that the relation between *K* and density could be described by the Gaussian curve

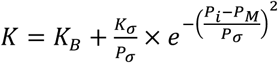

where *P*_*i*_ is the initial nematode density in nematodes per gram soil, *K*_*B*_ is the basal canopy size, *K*_*σ*_ is the normalized maximum canopy area that can be achieved over the *P*_*i*_ range, *P*_*σ*_ is the deviation around the nematode density allowing maximum growth, *P*_*M*_ is the nematode density at which maximum growth is achieved.

We found that the relation between *r* and density could be described by the exponential function

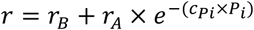

where *P*_*i*_ is the initial nematode density in nematodes per gram soil, *r*_*B*_ is the basal growth rate, *r*_*A*_ is the adaptive growth rate, and *c*_*Pi*_ is the translation constant, which translates the *P*_*i*_ to impact to growth rate.

Together, the function for *K* and *r* could be used to model the time series data of the entire experiment in the logistic growth model. We modelled the parameter values using *nls* and extracted confidence intervals using the *nlstools* package. In this model the tolerance limit can be defined as 2\**P*_*M*_ (where the achieved canopy area is equal to the achieved canopy area at *P*_*i*_ = 0).

## Results

### Density-response relationship between plant architecture and nematode inoculation density

The impact of biotic stresses on plant growth and development can be expressed as a function of inoculation density of causal agents of disease. To determine which plant growth parameters sensitively reflect the impact of nematode challenge on *A. thaliana*, we monitored changes in root system architecture and above-ground plant features at increasing inoculation densities of the beet cyst nematode *H. schachtii*. We first quantified different root system architecture components (*i*.*e*., primary root length, the number of secondary roots, and secondary root length) after inoculating *in vitro* cultured Arabidopsis plants with increasing densities (*P*_*i*_) of *H. schachtii* (Fig. 1a). At the system architecture level, we found that challenging Arabidopsis with *H. schachtii* at higher inoculation densities disturbs the typical regular patterning of emerging lateral roots (Fig. 1b). However, despite disturbing the lateral root patterning, the total root length did not change by increasing inoculation densities of *H. schachtii* (Fig. 1c). We did observe a decrease in primary root length at *P*_*i*_ of 5, and higher, but this was compensated by an increase in the number of secondary roots (Fig. 1c).

**Figure 1:**
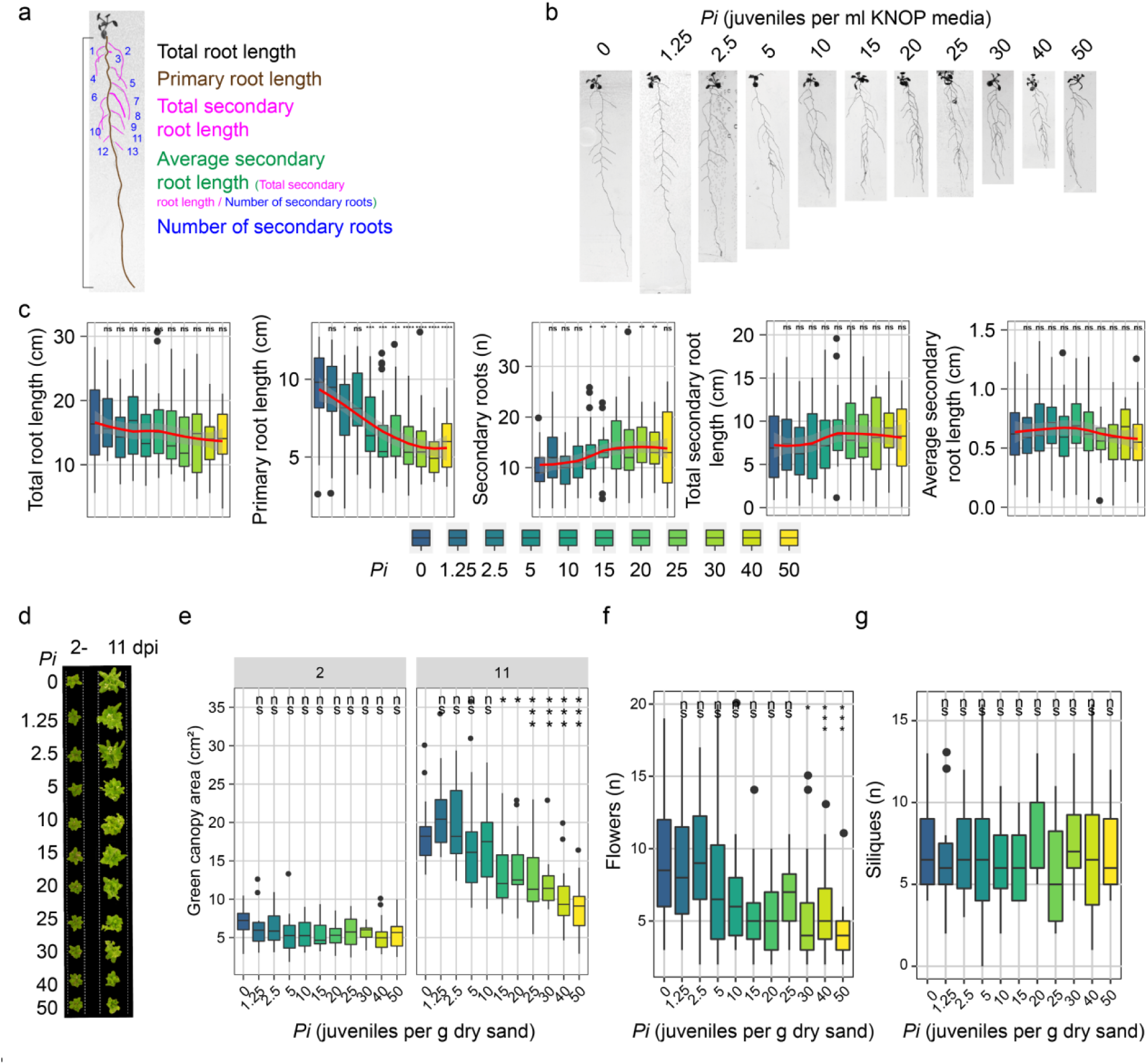
The primary root length and the green canopy area of Arabidopsis respond in a density-dependent manner to infection by *H. schachtii*. **a-c)** Nine-day-old Arabidopsis seedlings were inoculated with densities ranging from 0-50 of *H. schachtii* juveniles (ml modified KNOP media). The growth of the seedlings was monitored and measured at 7-days post-inoculation (dpi). **a)** Overview of root system architecture components that we have quantified. **b)** Representative images of Arabidopsis root system at 7 dpi. **c)** Total root length (cm), primary root length (cm), total secondary root length (cm), average secondary root length (cm), and the number of secondary roots (n). Data was analysed by one-way ANOVA with post-hoc Tukey HSD; ns= not significant, *p< 0.05, **p< 0.01, ***p<0.001 (n=24-30). **d-g)** Twenty-one-day-old Arabidopsis seedlings cultivated in soil were inoculated with increasing densities of second juveniles of *H. schachtii* (from 0 to 50 juveniles per g dry sand). **d)** Representative images of Arabidopsis seedlings aboveground at 2 and 11 dpi. **e)** Green canopy area (cm^2^) at 2 and 11-dpi with increasing densities (Pi). **f)** Number of flowers at 11 dpi. **g)** Number of siliques at 11 dpi. Data was analysed by one-way ANOVA with post-hoc Tukey HSD; ns= not significant, *p< 0.05, **p< 0.01, ***p<0.001 (n=16-20).

Next, we monitored the impact of *H. schachtii* densities on aboveground plant growth and development of *A. thaliana* cultivated in soil by daily assessing changes in green canopy area, the number of flowers, and the number of siliques (Fig. 1d). The above-ground plant growth/development over time showed clear detrimental effects linked to increased *H. schachtii* infection pressure (Fig. S2a). At 11 days post inoculation (dpi), we observed the biggest differences in green canopy area between inoculation densities, including a tolerance limit of Arabidopsis for *H. schachtii* at 10 juveniles per gram soil (Fig. 1d and e). We also counted the number of flowers and siliques over time (Fig. S2b and c). Likewise, we also observed a density-dependent response for the number of flowers, albeit at higher inoculation densities (Fig. 1f). By contrast, we did not find a significant response in the number of siliques by inoculation density (Fig. 1g). In conclusion, both primary root length and green canopy area can be used to assess the impact of nematode-induced biotic stress on *A. thaliana*.

We tested whether the green canopy area could also be used as a proxy for belowground adaptations to nematode challenge, as repeated measurements on root system architecture requires a more artificial experimental design. To address this question, we calculated the Spearman’s Rank Order Correlation Coefficient for our measurements of green canopy area and root system architecture components acquired through different experimental designs (Fig. S3). Indeed, we found that green canopy area strongly correlates with primary root length (*R*_*2*_ = 0.89, *P* <0.0001) (Fig. S3b). Green canopy area also anticorrelated well with the number of secondary roots (*R*_*2*_ *=* −0.63, *p* = 0; Fig. S3e) and total secondary root length (*R*_*2*_ = −0.64, *p* = 0.04; Fig. S3c). As expected, we found no significant correlation between the green canopy area and average secondary root length (Fig. S3d) and total root length (Fig. S3a). Based on this analysis, we focused further on developing a high-throughput automated phenotyping platform using the green canopy area of Arabidopsis cultivated in soil to assess the impact of *H. schachtii* at different inoculation densities over time.

### High-throughput monitoring of density-response relationships between nematode densities and green canopy area

Our high-throughput automated monitoring platform can record and analyse changes in green canopy area of 960 Arabidopsis plants simultaneously for a period of twenty-one days (Fig. 2a). The platform design includes stainless steel frames that can hold 200 mL pots filled with soil (Fig. S1). The frames fix the pots in position and give anchor to perforated matte black cover plates which maximise the contrast of the green canopy area. The frames and cover plates match the size of a flooding table placed in a fully automated climate room. Each frame contains 160 perforated holes to sow seeds and inoculate the pots. Photographs can be made at any desired time interval by high-resolution cameras mounted at fixed positions above the cover plates. To monitor the impact of nematode challenge on green canopy area, photographs were taken at one-hour intervals for 21 days during daytime (i.e., 15 pictures per day). To collect data on individual plants, the pictures were processed by colour channel decomposition in Adobe Photoshop and analysed in ImageJ (Fig. 2a). With every experiment, this results in 302,400 pictures of individual plants (315 pictures per plant).

**Figure 2:**
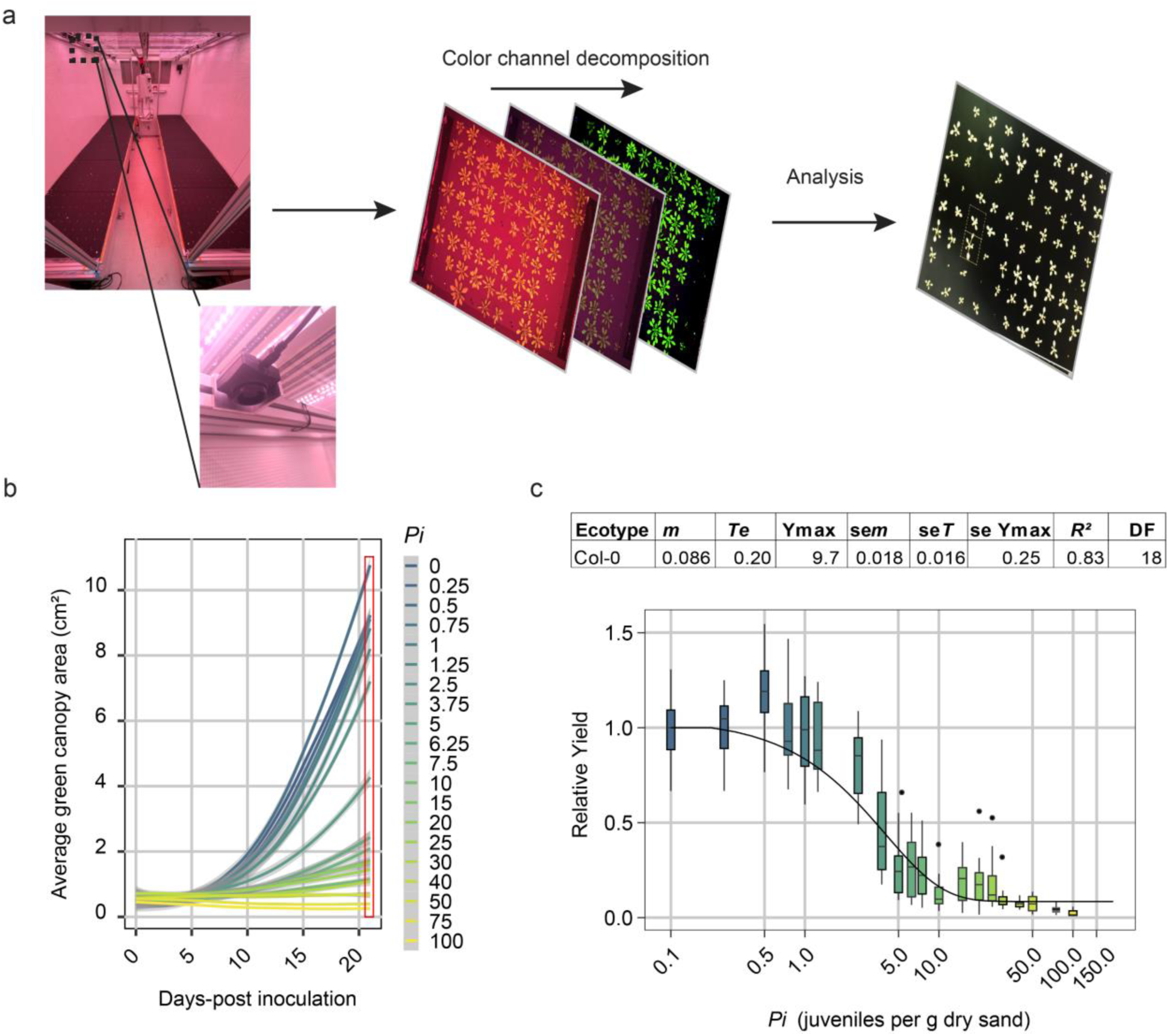
The relationship between the inoculation density (*Pi*) of *H. schachtii* and green canopy area (Relative Yield). Nine-day-old Arabidopsis seedlings were inoculated with 20 densities of *H. schachtii* juveniles (0 to 100 J2s per g dry sand) in 200 mL pots containing 200 grams of dry sand. **a)** Experimental setup and colour channel decomposition of photos to extract the green leaf surface from the images. **b)** Average growth curve of Arabidopsis plants inoculated with different inoculum densities of *H. schachtii* from 0-21 dpi. Line fitting was based on a LOESS regression. Red box indicates the data that is used for fitting the Seinhorst yield loss equation. **c)** Fitting according to the Seinhorst yield loss equation. Parameter values for Seinhorst’s Eq. for the relation between initial population density (*Pi*) of *H. schachtii* and measured leaf surface area. *Pi* and tolerance limit (*Te*) are expressed in *H. schachtii* (g dry sand)^-1^ while, the minimal yield (*m*) is the lowest proportion of the maximum green canopy area (cm^2^) (*Ymax*) at 21dpi. The goodness of the fit of the model on the data is expressed as the coefficient of determination (*R*^*2*^).

To validate our system, we inoculated 480 *A. thaliana* Col-0 seedlings with increasing densities of *H. schachtii* (*P*_*i*_ 0 to 100 juveniles per g dry sand) and monitored their growth for a period of three weeks. The green canopy area rendered from the recordings was collected and analysed (Movie 1; Fig. 2b). We confirmed that increasing densities of *H. schachtii* decreased the growth of Arabidopsis plants. To test if changes in the green canopy area fit the SYLM, we tested our green canopy area at 21 dpi for its fit (equation 1) (Seinhorst, 1985). We found the typical inverted sigmoid curve and that our data fits well to the Seinhorst Yield Loss Model (*R*_*2*_ = 0.83; Fig. 2c), indicating that there is a strong relationship between nematode densities and the green canopy area. Also, we found that the tolerance limit (*Te*) is placed around *P*_*i*_ 0.21 (Fig. 3c). Altogether, we concluded have developed a high-throughput system in which we can quantify the relation between green canopy size and cyst nematode inoculum densities using the Seinhorst equation for yield losses for Arabidopsis.

**Figure 3:**
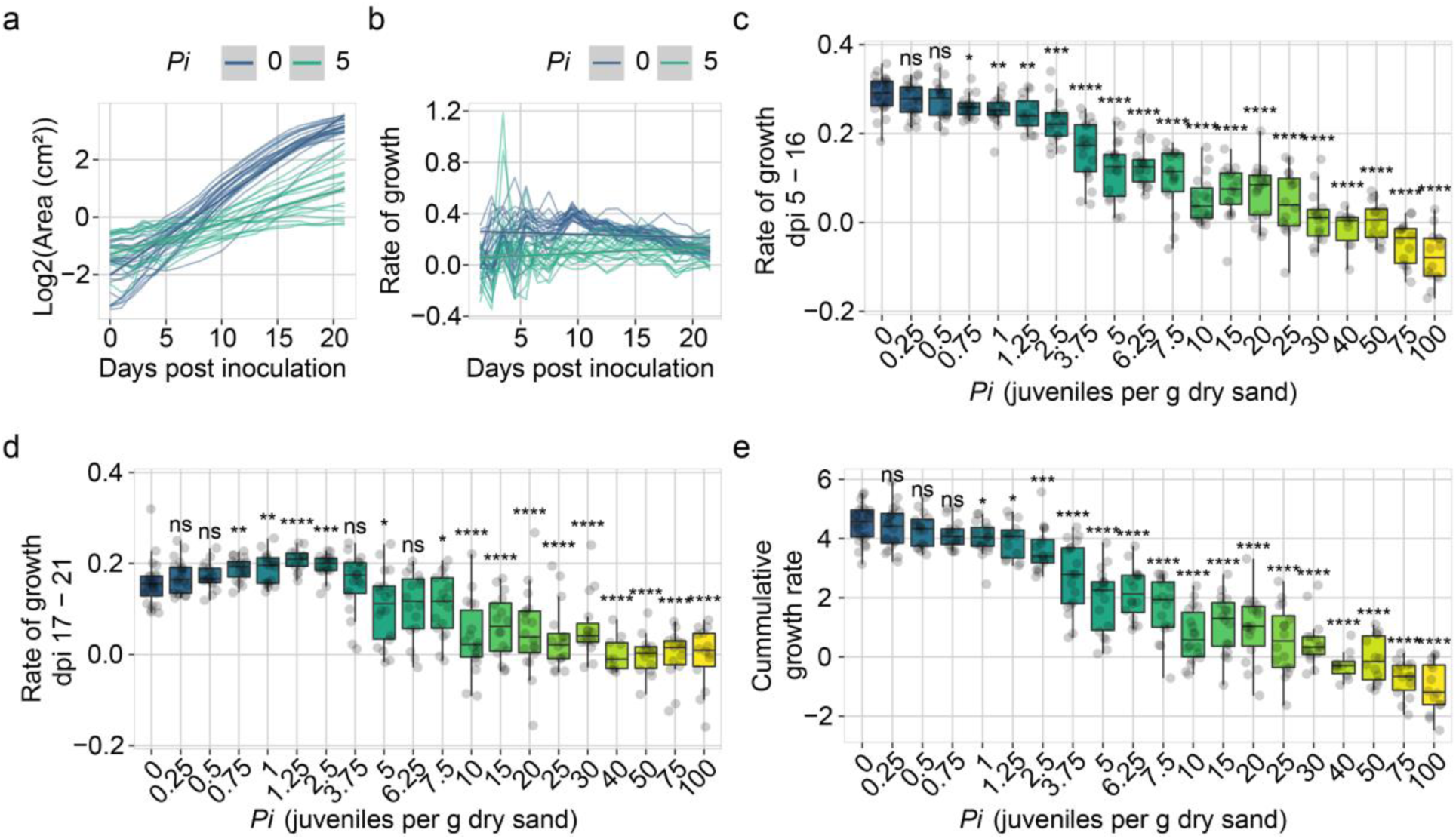
Quantification of tolerance limit of Col-0 to *Heterodera schachtii* based on growth rate. Nine-day old Arabidopsis seedlings were inoculated with 20 densities (*Pi*) of *H. schachtii* juveniles (0 to 100 juveniles per g dry sand). **a)** Green canopy area of plants treated with *Pi* 0 (blue line) or 5 (aquamarine line) from 0 to 21 days-post inoculation (dpi). **b)** Rate of growth of plants treated with *Pi* 0 (blue line) or 5 (aquamarine line) from 0 to 21 dpi was calculated using equation 2. **c)** Median rate of growth of plants inoculated with increasing *Pi’s* growing from 5 to 16 dpi. **d)** Rate of growth (cm^2^) of plants inoculated with increasing *Pi’s* growing from 17 to 21 dpi. **e)** Cumulative growth rate of plants inoculated with increasing *Pi’s*. **c-e)** Dots represent individual plants. Data was analysed with a Wilcoxon Rank Sum test; ns= not significant, *p< 0.05, **p< 0.01, ***p<0.001 (n=10-24).

### Quantifying tolerance limits on growth rates

The SYLM does not accommodate for compensation effects we observed at low inoculation densities (Fig. 2c). Based on the boxplots, the tolerance limit *Te* is between *P*_*i*_*s* 1.25 and 2.5, whereas the SYLM placed it at 0.21. Another drawback of the SYLM is that data of a single timepoint of a series is tested for fit, ignoring changes over time. To take the dynamics in plant response into account, we sought an alternative way to quantify tolerance limits that uses all data collected over time (Fig. 2b). It is described that a delay in plant growth occurs at lower nematode densities and that higher nematode densities may lead to earlier cessation of growth (Seinhorst, 1985). Therefore, we extracted the growth rate (Equation 2) of plants inoculated with different nematode densities from 0 to 21 dpi to test if changes in growth rates can be used to determine tolerance limits.

First, we calculated the growth rate per day of individual plants that were inoculated with increasing *H. schachtii* densities (Fig. S4, S5, S6; Fig. 3a, b). Plants that were mock-inoculated showed a consistent growth rate in green canopy area between 5 and 15 dpi (Fig. 3c). Plants inoculated with *P*_*i*_ 0.75 or higher had a lower growth rate compared to mock-treated plants (Fig. 3c). From 15 dpi onwards, growth rates of mock-inoculated plants reached a stationary phase, while plants inoculated with nematodes (*P*_*i*_ 0.75 to 2.5) continued their exponential growth rate (Fig. 3d). Finally, we calculated the cumulative growth rate, which is the sum of all growth rates over time per plant (Fig. 3e), which showed that plants treated with *P*_*i*_ 1 or higher grew significantly slower compared to mock-inoculated plants. Based on these results, we conclude that the tolerance limit of Arabidopsis Col-0 to for *H. schachtii* is around *P*_*i*_ *=* 1.

### Quantifying tolerance limit to *M. incognita*

Root-knot nematodes migrate through the roots intercellularly and therefore cause less damage than cyst nematodes. Hence, we hypothesized that the tolerance limit for *M. incognita* infections should be at a higher *P*_*i*_ than for *H. schachtii*. To this end, 320 nine-day old Arabidopsis seedlings were inoculated with 18 different densities (*P*_*i*_ 0 to 100) of *M. incognita* and monitored for 21 days. We fitted the data to the SYLM and found a *Te* of 0.57 (Fig. S7). Like we observed for *H. schachtii*-infected plants, we found that the model underestimates the *Te*. We thus also calculated the growth rates to capture the dynamics of the system (Fig. S8, Fig. 4a-e). Here, we observed that the growth rates of plants between 5 and 16 days-post inoculation were less affected by *M. incognita* than by *H. schachtii* (Fig. 4c). The growth rates of plants treated with *P*_*i*_ 1.25 or higher were significantly slower. From 17 to 21 dpi, we observed that plants inoculated with *M. incognita* at *P*_*i*_ 1 - 7.5 tended to have a higher growth rate than mock-inoculated plants (Fig. 4d). Plants inoculated with *P*_*i*_ 40 and 100 stopped growing altogether. Next, we calculated the cumulative growth rate, which showed that plants treated with *P*_*i*_ 2.5 or higher grow significantly slower compared to mock inoculated plants (Fig. 4e). Based on these results, we concluded that our high-throughput phenotyping system can also determine tolerance limits to root-knot nematodes and that the tolerance limit of *A. thaliana* Col-0 to *M. incognita* infection is around *P*_*i*_ 2.5.

**Figure 4:**
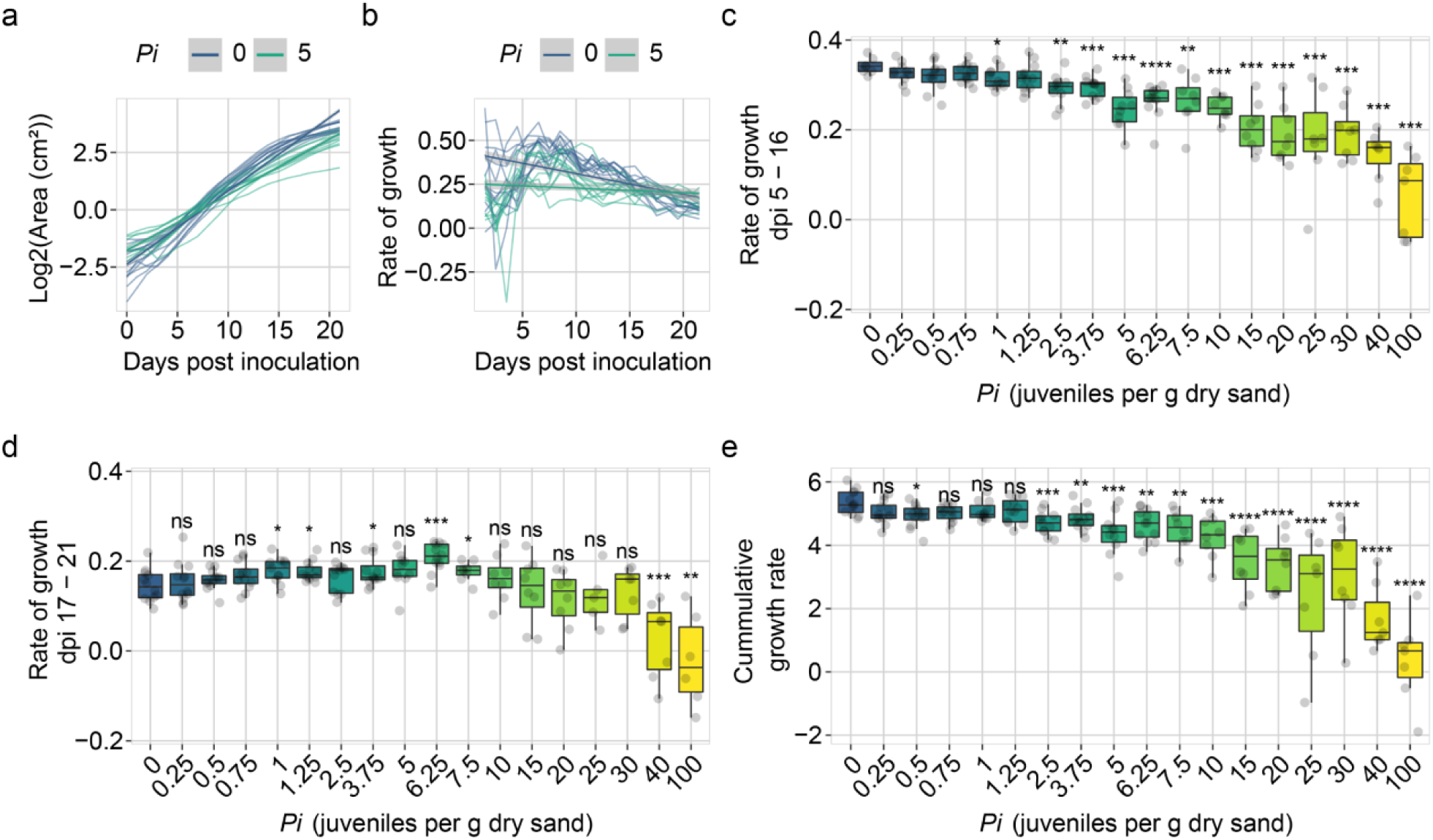
Quantification of tolerance limit of Col-0 to *Meloidogyne incognita* based on growth rate. Nine-day-old Arabidopsis seedlings were inoculated with 18 densities (*Pi*) of *M. incognita* juveniles (0 to 100 juveniles per g dry sand. **a)** Green canopy area of plants treated with *Pi* 0 (blue line) or 5 (aquamarine line) from 0 to 21 days-post inoculation (dpi). **b)** Rate of growth of plants treated with *Pi* 0 (blue line) or 5 (aquamarine line) from 0 to 21 dpi was calculated using equation 2. **c)** Rate of growth of plants inoculated with increasing *Pi’s* growing from 5 to 16 dpi. **d)** Rate of growth of plants inoculated with increasing *Pi’s* growing from 17 to 21 dpi. **e)** Cumulative growth rate of plants inoculated with increasing *Pi’s*. **c-e)** Dots represent individual plants. Data was analysed with a Wilcoxon Rank Sum test. ns= not significant, *p< 0.05, **p< 0.01, ***p<0.001 (n=10-12).

### Arabidopsis Col-0 is more tolerant to *M. incognita* than to *H. schachtii*

Next, we correlated the growth rates of *M. incognita* inoculated plants to *H. schachtii* inoculated plants per timepoint and per inoculation density to determine if the growth rates of *A. thaliana* Col-0 are less affected by initial nematode densities of *M. incognita* than by *H. schachtii* (Fig. S9a and b). Both correlations based on timepoint and inoculum density between *H. schachtii* and *M. incognita* inoculated plants indicate that plants inoculated with *M. incognita* retain higher growth rates than plants inoculated with the same density of *H. schachtii*. Subsequently, we tested if there are significant differences between growth rates per *P*_*i*_ and per timepoint. For instance, we observed that *H. schachtii* infection has more impact on *A. thaliana* growth rates than *M. incognita* at *P*_*i*_ 5, especially from 5 to 10 dpi (Fig. S10a). From 11 to 21 dpi, the differences were smaller but often still significant. Then, we calculated the difference in growth rate by extracting the *H. schachtii* treated growth rates from the *M. incognita* treated growth rates per timepoint and *P*_*i*_ (Fig. S10b). We found that most of the growth rates of plants inoculated with *M. incognita* are higher than those of *H. schachtii* inoculated plants. Also, the largest differences in growth rates were visible from 5 to 10 dpi (Fig. S10b and c).

Pair-wise comparisons of green canopy areas between different treatments are cumbersome and drawing conclusions remain elusive. Therefore, we fitted and tested our growth dynamics of green canopy areas (Fig. 2, S7, 3, and 4) to a logistic growth model to determine if Arabidopsis Col-0 is more tolerant to *M. incognita* and *H. schachtii*. The logistic growth model determined the maximum canopy area and the intrinsic rate of growth. We found that the maximum canopy area *K* had relation with *P*_*i*_ that could be fitted with a Gaussian curve (Fig. 5a and c) and that the relation between the intrinsic growth rate *r* and *P*_*i*_ could be fitted with a hyperbolic curve (Fig. 5b and d). These fits allowed us to estimate the tolerance limits, which were *P*_*i*_ = 3.3 (95% CI: 3.0-3.7) for *H. schachtii* and 5.5 (95% CI: 4.6-6.4) for *M. incognita*. We therefore conclude that *A. thaliana* Col-0 is more tolerant to biotic stress by *M. incognita* infection than *H. schachtii*.

**Figure 5:**
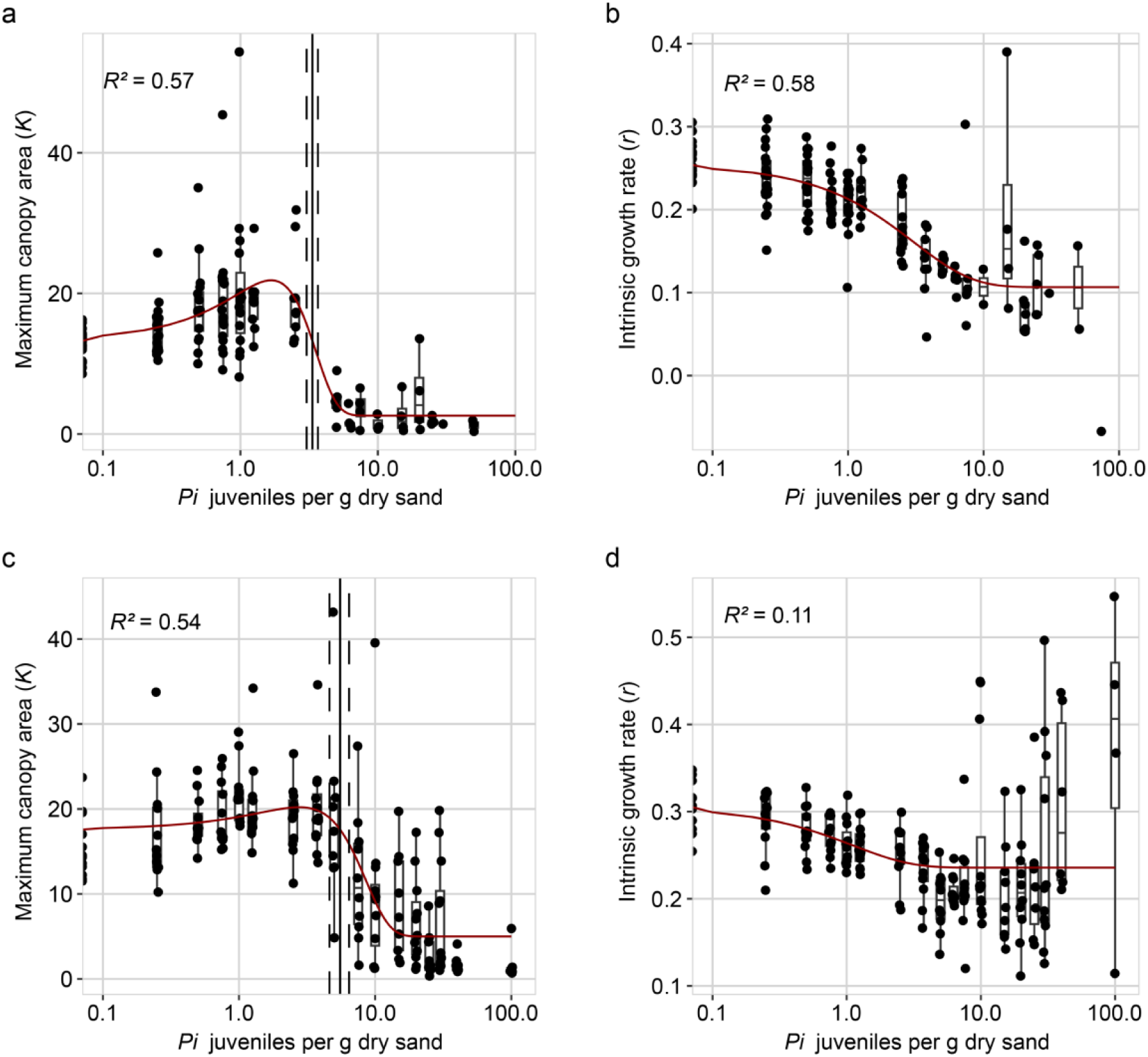
*Arabidopsis thaliana* Col-0 is more tolerant to *Meloidogyne incognita* than *Heterodera schachtii*. Fitted logistic growth rate parameters of *Arabidopsis thaliana* plants infected with *H. schachtii* or *M. incognita*. (**A**) The maximum canopy area *K* per infection density of *H. schachtii*. The fitted line is from a Gaussian curve, the *R*^*2*^ is from the fitted curve. (**B**) The intrinsic growth rate *r* per infection density of *H. schachtii*. The fitted line is from an exponential model, the *R*^*2*^ is from the fitted curve. (**C**) Like (**A**) but for infections with *M. incognita*. (**D**) like (**B**) but for infections with *M. incognita*. Solid line indicates the tolerance limit. Dashed line indicates the confidence interval.

## Discussion

Plants show intraspecific variation in how much damage by root-parasitic nematodes they can cope with before growth delays or ceases altogether (Potter & Dale, 1994). Unfortunately, uncovering the underlying molecular mechanisms of tolerance remains challenging and laborious since no tractable genetic model system is available. The goal of this research was twofold: (i) assess the suitability of *A. thaliana* as model for tolerance to root-parasitic nematodes and (ii) develop a high-throughput phenotyping platform to quantify and measure tolerance levels. The suitability of *A. thaliana* to measure tolerance was demonstrated with the beet cyst nematode *H. schachtii* and the root-knot nematode *M. incognita*. We present a high-throughput phenotyping platform that allows for the collection of continuous plant-growth data (here with a resolution of one data point per hour). Continuous data is an important improvement upon the end-point data which is normally used to determine tolerance levels of plants to nematode infection. Based on our findings, we believe that our phenotyping platform will enable us to uncover the underlying genetic mechanisms of tolerance.

We show that the green canopy area in *A. thaliana* is a suitable proxy to study tolerance limits and growth responses of plants to root-parasitic nematode infections. The nematodes *H. schachtii* and *M. incognita* infect root tissues, but it is difficult and laborious to follow and quantify root development over time. Although we measured individual root traits, the throughput and therefore the data density were low. Previously, it was shown that green canopies are suitable to monitor the effect of nematode infection on sugar beet (Joalland *et al*., 2016, 2017). Therefore, we monitored green canopy area growth of individual plants in a non-destructive way by taking pictures. We showed that green canopy area is a good proxy for below-ground responses to nematode infection. Multiple root traits correlated well with the green canopy area. Furthermore, our approach allows to capture changes in growth rates in response to nematode infection over time. This allowed us to identify dynamic damage responses and compensatory growth rates of nematode-infected Arabidopsis plants.

We observed three distinct effects of infections by plant parasitic nematodes on plant growth-rate: an initial damage response, a growth rate-recovery at early timepoints, and a growth compensation response at later timepoints. First, we observed that plants showed an initial damage response by slowing growth rates shortly after *M. incognita* inoculation. The growth rates recovered four days later. Similar observations were made in root growth upon prolonged periods of invasion by the cyst nematode *Heterodera avenae* (Rawsthorne & Hague, 1986). Secondly, we observed that *H. schachtii* inoculated plants grew significantly slower compared to *M. incognita* inoculated plants at 5 to 10 dpi under similar nematode inoculation densities. It is unclear whether the differential growth rates are due to differences in pathogenicity between the two nematode species, the types of damage that these two nematodes cause, or by the interaction of the nematode species and Arabidopsis. Thirdly, at the later timepoints (16-21 dpi), plants inoculated with lower dosages of *H. schachtii* and *M. incognita* showed increased growth-rates compared to mock-inoculated plants. To our knowledge, this is the first observation of – next to growth-recovery at early timepoints – growth-compensation at later stages of infection. Yield losses by nematodes could be minimized if characteristics like growth recovery and compensation are incorporated in breeding programs. However, this requires more insight in the genetic (in)dependence of these growth-response traits.

There seems to be a link between the infection stages of the nematodes and the changes that we measured in growth-kinetics of host plants. The nematode infection process starts with root invasion, followed by migration through root tissue, after which feeding sites are formed leading to loss of assimilates via these feeding sites. First, we believe that the slower growth rate during the first four/five days is due to *M. incognita* and *H. schachtii* invasion and migration. A faster recovery response by the plant to *M. incognita* infection than to *H. schachtii* infection could have three reasons: 1) *M. incognita* needs less time to induce a feeding site and therefore the plant can continue growth faster, 2) *M. incognita* causes less stress due to its different way of invasion and migration, or 3) the feeding site is induced at a different place in the root. Specifically, cyst nematodes can invade the root at different locations along the root axis, and subsequently migrate intracellularly through the cortex towards the vascular cylinder. Whereas root-knot nematodes invade the root at the root tip and migrate intercellularly through the cortex and the apical meristem to enter the vascular cylinder from below. (Wyss & Zunke, 1986; Wyss, 1992; Wyss *et al*., 1992; Kyndt *et al*., 2013). Finally, we measured the loss of assimilates by nematodes resulting in slower plant growth. After partial growth recovery in response to lower nematode densities for *M. incognita* and *H. schachtii* treated plants, we observed that growth rates slowed down again around 7 dpi. Mechanical or physiological damage during feeding site initiation and expansion, and consequently withdrawal of plant assimilates are important factors that contribute to reducing plant growth and delaying development (Trudgill, 1991). Being able to measure the effects of different nematode life stages (invasion/migration and feeding) on plant growth could give new and interesting insights in how plants respond to different types of stress (i.e. mechanical damage versus loss of assimilates). Given we are able to phenotypically discriminate growth responses to different types of stresses indicates that tolerance is a complex trait. We hypothesise that tolerance is a combination of multiple (simultaneous) processes with each their own genetic architecture.

Our observation of the compensatory growth response shows that there are two limitations in the application of the SYLM on nematode-infected *A. thaliana*. First, the SYLM underestimates the tolerance limits *Te* for *H. schachtii* and *M. incognita* infections in Arabidopsis. Yet, the overall density response in our data fitted the model well. Similar observations were made previously where the *Te* strongly deviated between biological replicates (Teklu *et al*., 2022). We conclude that although the SYLM does not perform well at low-densities, it is suitable to predict minimal yields. Second, the SYLM is inefficient as it uses only end-point data (Seinhorst, 1985; Norshie *et al*., 2011; Sasanelli *et al*., 2013; Been *et al*., 2015; Moosavi, 2015), like tuber yield or shoot biomass. The SYLM does not capture the complex growth dynamics. Insight in the complex growth dynamics is necessary to separate different tolerance mechanisms.

To conclude, plants display intraspecific growth responses to nematode infection. These differences in tolerance levels are important agronomic trait of crops as they determine the level of stress that the plant can handle before growth is delayed. Due to the lack of a tractable genetic model system, the molecular mechanisms underlying tolerance remain elusive. Our high-throughput phenotyping platform is capable of monitoring the effect of nematode infection on the growth of the model plant *A. thaliana* will provide novel insights into how plants mitigate the impact of damage by nematodes and how plants try to compensate growth delays caused by the loss of assimilate by feeding nematodes. Specifically, it will provide novel insights in the molecular and genetic mechanisms that underlies tolerant growth responses, which provide to an additional layer of protection for securing global food production besides disease resistance.

## Abbreviations

J2s: Infective juveniles
SYLM: Seinhorst Yield Loss Model
*Te*: Tolerance limit
*P*^*i*^: Nematode density

## Supporting Information

Additional Supporting Information may be found in the online version of this article

## Author contributions

JJW, GS and MGS conceived the project. JJW and LD designed the high-throughput platform experiment. JJMS designed automated data collection software. JJW, DS, and CCS collected data. MGT provided scripts for SYLM analysis. Data analysis was designed by MGS with help from JJMS and data was analyzed and interpreted by JJW and MGS. JJW, GS, and MGS wrote the article. JLLT and AG provided critical feedback on the manuscript. All co-authors provided input for the submitted version.

## Data availability

All relevant data can be found within the manuscript, Github (https://git.wur.nl/published_papers/willig_2023_camera-setup), or protocols.io (DOI: dx.doi.org/10.17504/protocols.io.kqdg39167g25/v1) (Private link for reviewers: https://www.protocols.io/private/C82F27F2BE7511ED91B00A58A9FEAC02 to be removed before publication.)

## Supplementary Information

**Figure S1:**
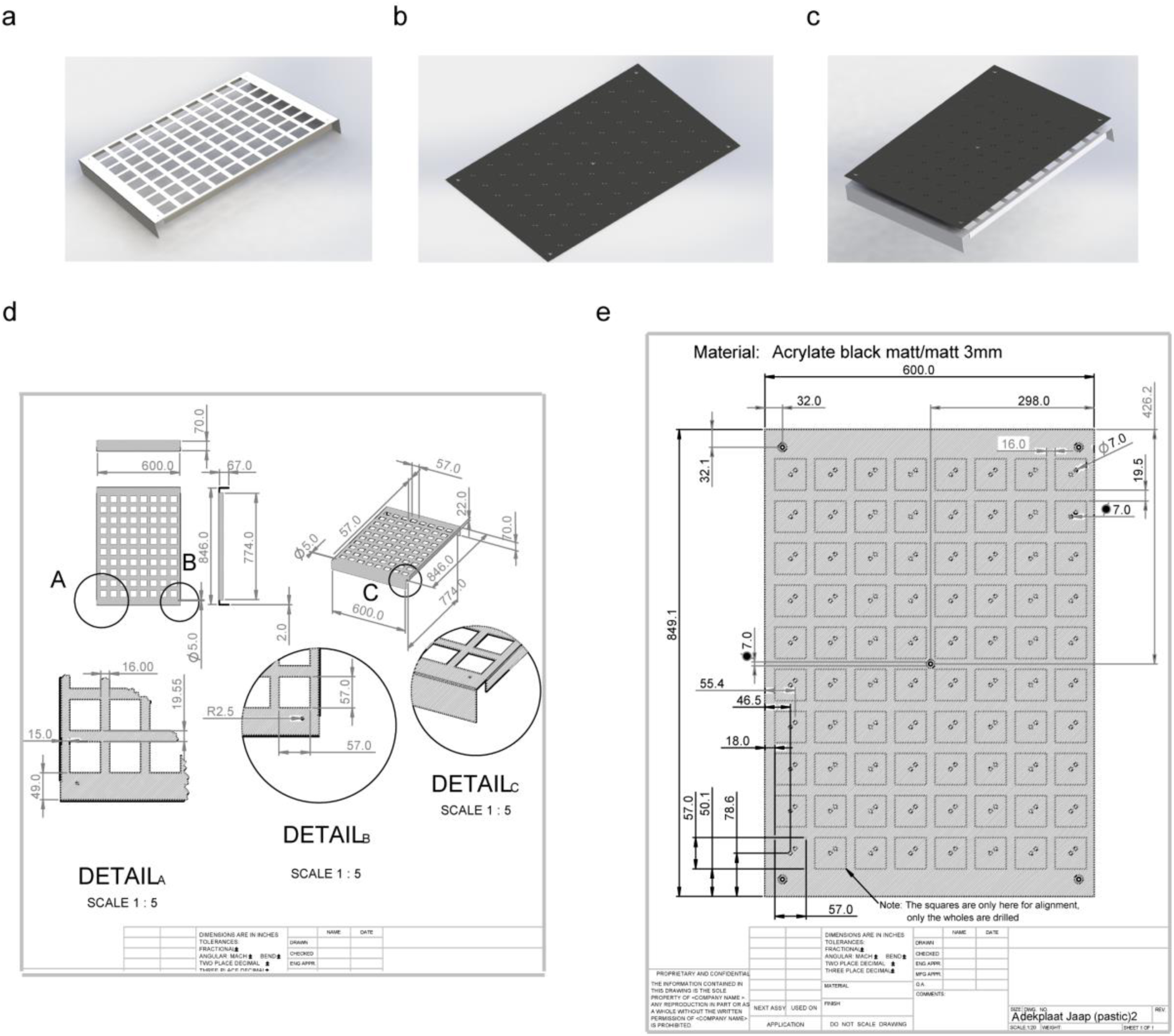
Overview of experimental setup. **a and d)** Stainless steel frames that we designed for holding pots. **b and e)** 3 mm thick black nonreflective foamed PVC coversheet drilled with countersunk holes and holes to inoculate nematodes. **c)** Black plates are attached to the aluminium frames with screws.

**Figure S2:**
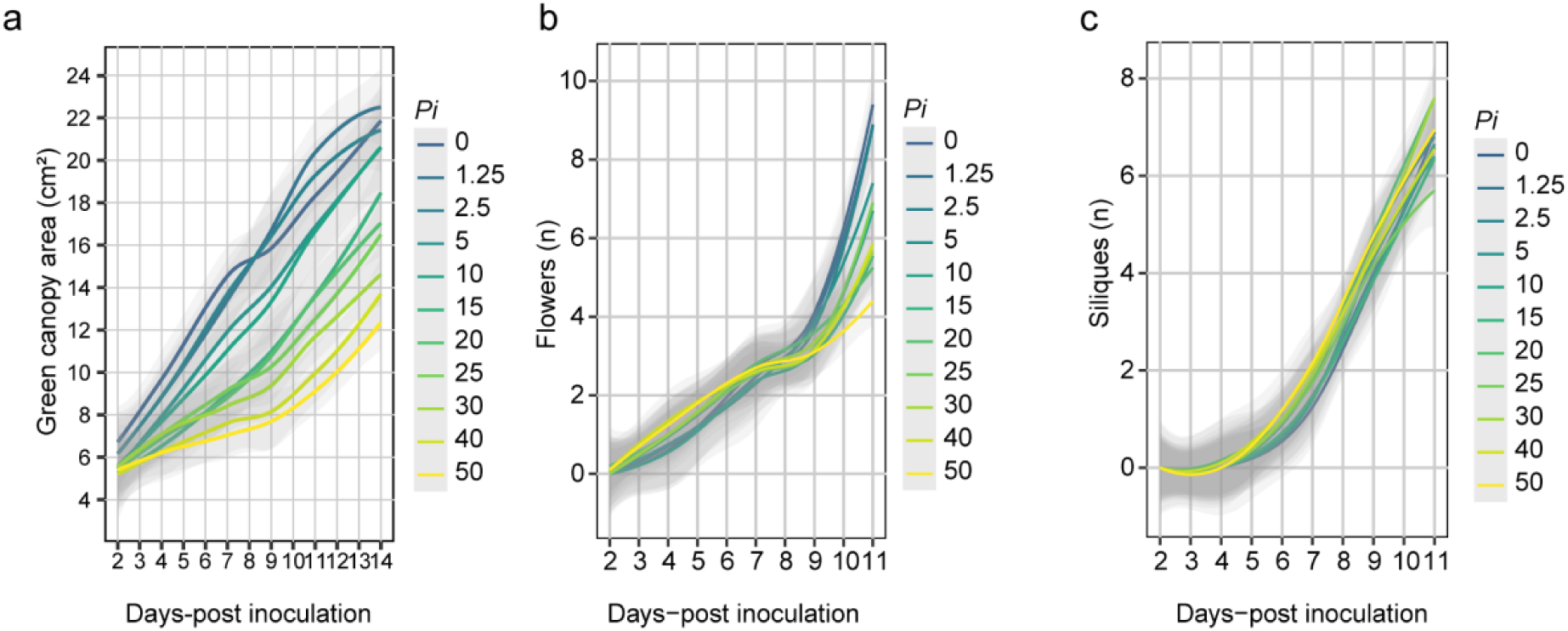
The primary root length and the green canopy area of Arabidopsis responds in a density-dependent manner to infection by *Heterodera schachtii*. Twenty-one-days-old Col-0 seedlings were inoculated with increasing densities of second-stage juveniles of *H. schachtii* (0-50 juveniles per g dry sand). At different timepoints, we counted the number of flowers and siliques and at 28 dpi we counted the number of cysts. **a)** Effect of nematode inoculations on the green canopy over time. **b)** Effect of nematode inoculations on the number of flowers over time. **c)** Effect of nematode inoculations on the number of siliques over time.

**Figure S3:**
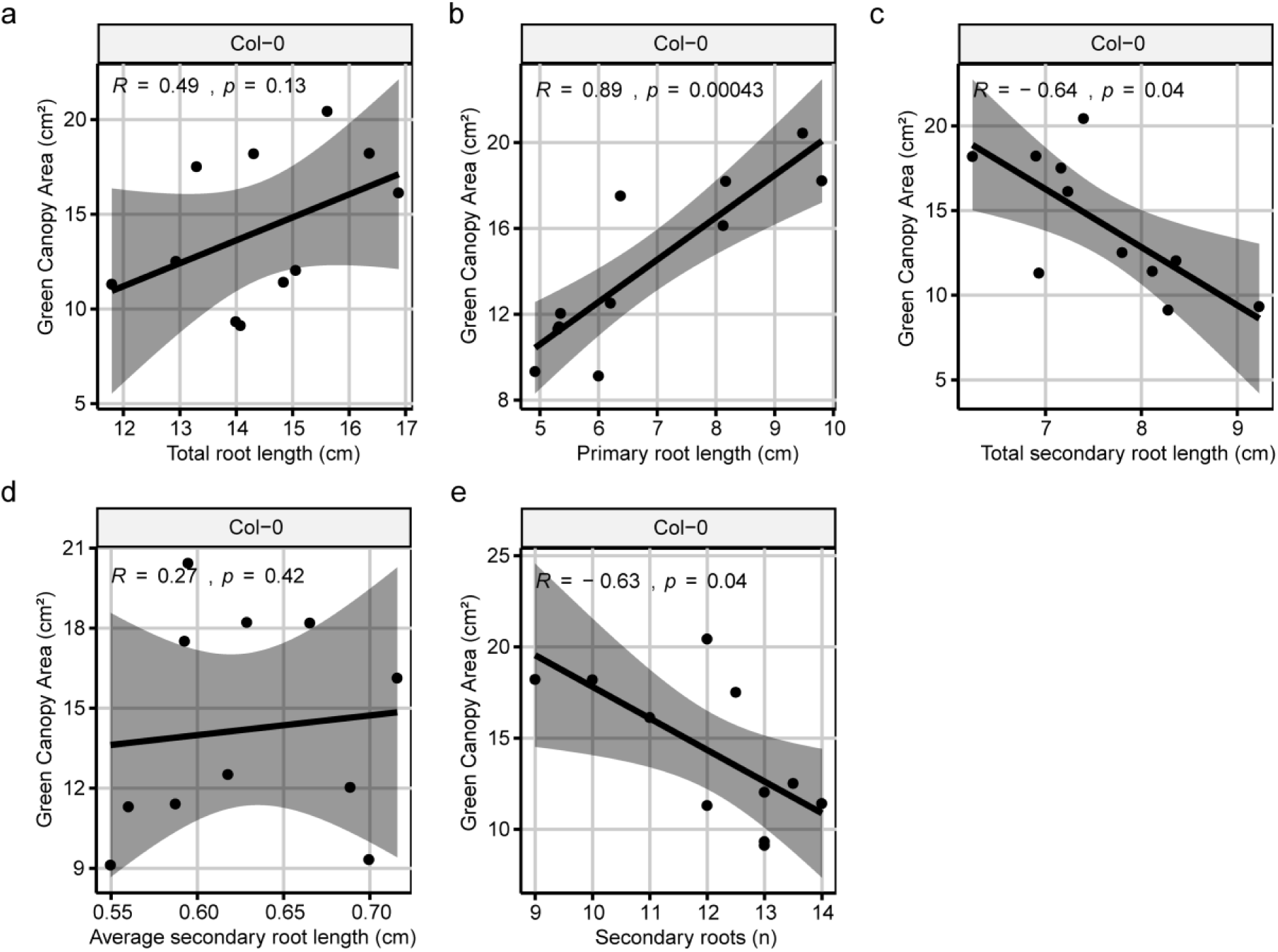
Spearman correlation coefficients calculated between the average of different measurements of root components at seven-days post inoculation and green canopy area at eleven-days post-inoculation. Spearman correlation coefficients were calculated on average values of 16-20 replicates of green canopy measurements and 24-30 replicates of root measurements using R software. **a)** Correlation between green canopy area and total root length. **b)** Correlation between green canopy area and primary root length. **c)** Correlation between green canopy area and total secondary root length. **d)** Correlation between green canopy area and average secondary root length. **e)** Correlation between green canopy area and the number of secondary roots.

**Figure S4:**
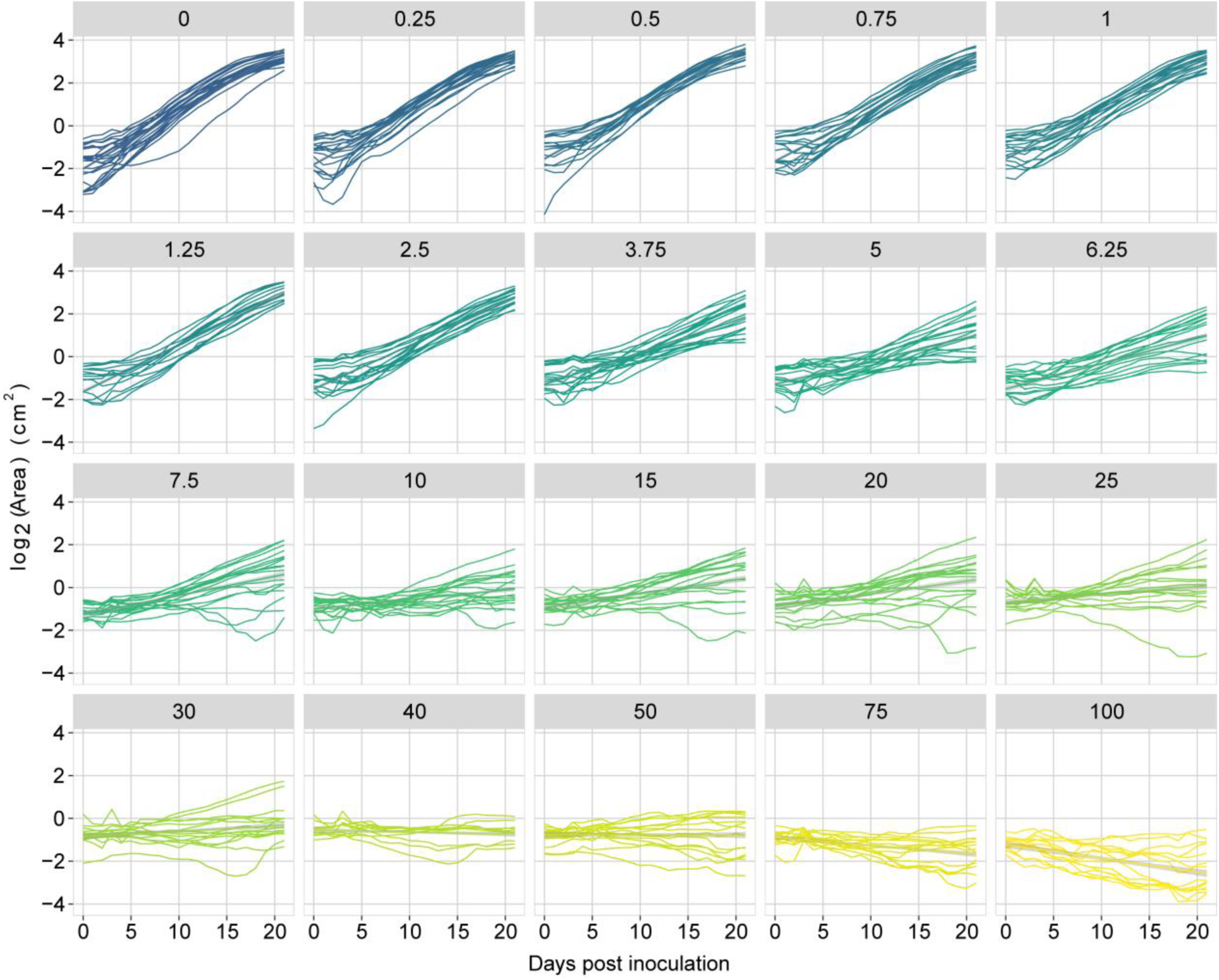
Growth of Arabidopsis Col-0 plants inoculated with increasing densities of *Heterodera schachtii*. Nine-day-old Arabidopsis seedlings were inoculated with 20 densities (*Pi*) of *H. schachtii* juveniles (0 to 100 juveniles per g dry sand). Lines represent the growth of individual plants.

**Figure S5:**
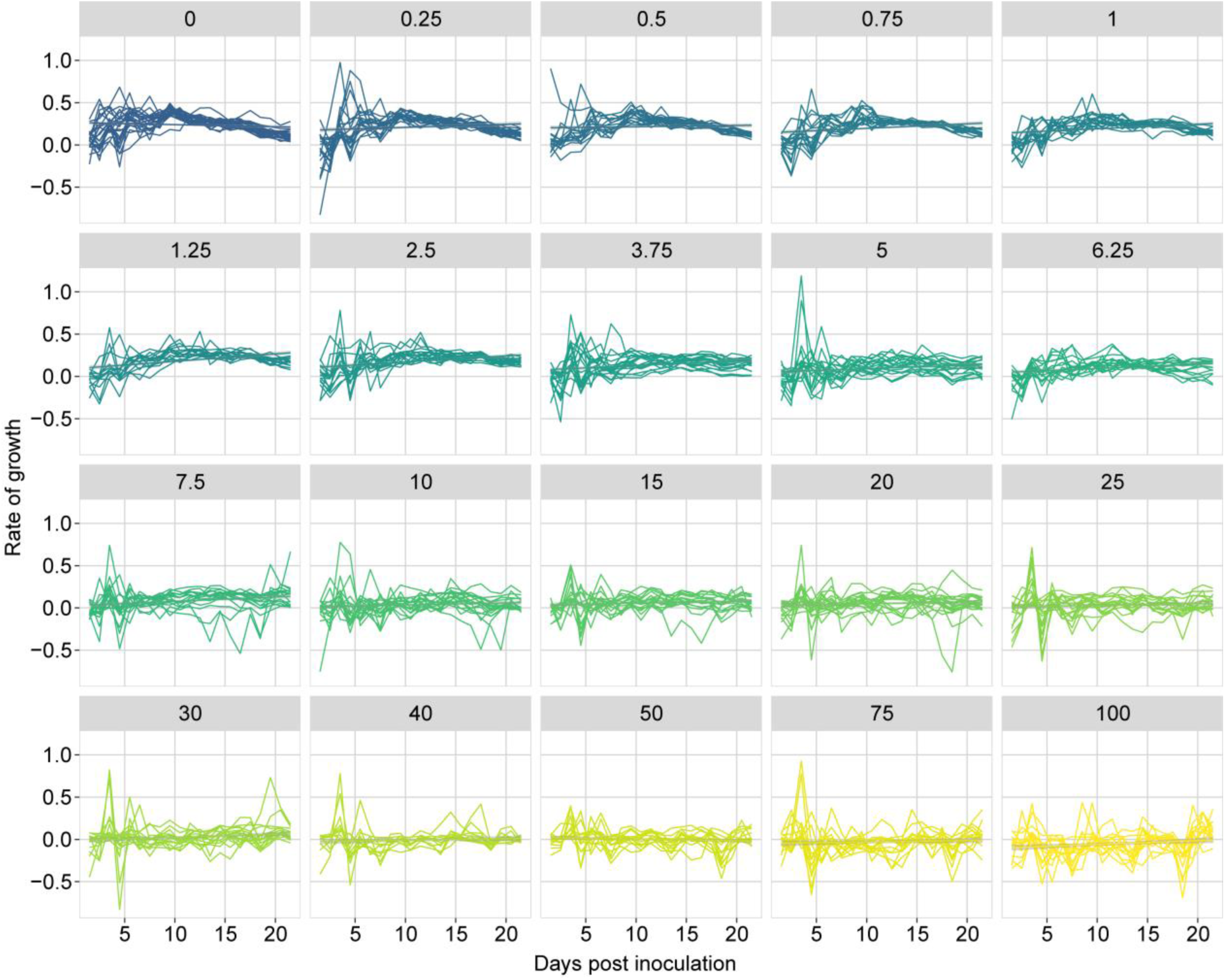
Growth rate of Arabidopsis Col-0 plants inoculated with increasing densities of *Heterodera schachtii*. Nine-day-old Arabidopsis seedlings were inoculated with 20 densities (*Pi*) of *H. schachtii* juveniles (0 to 100 juveniles per g dry sand). Lines represent the growth rates of individual plants.

**Figure S6:**
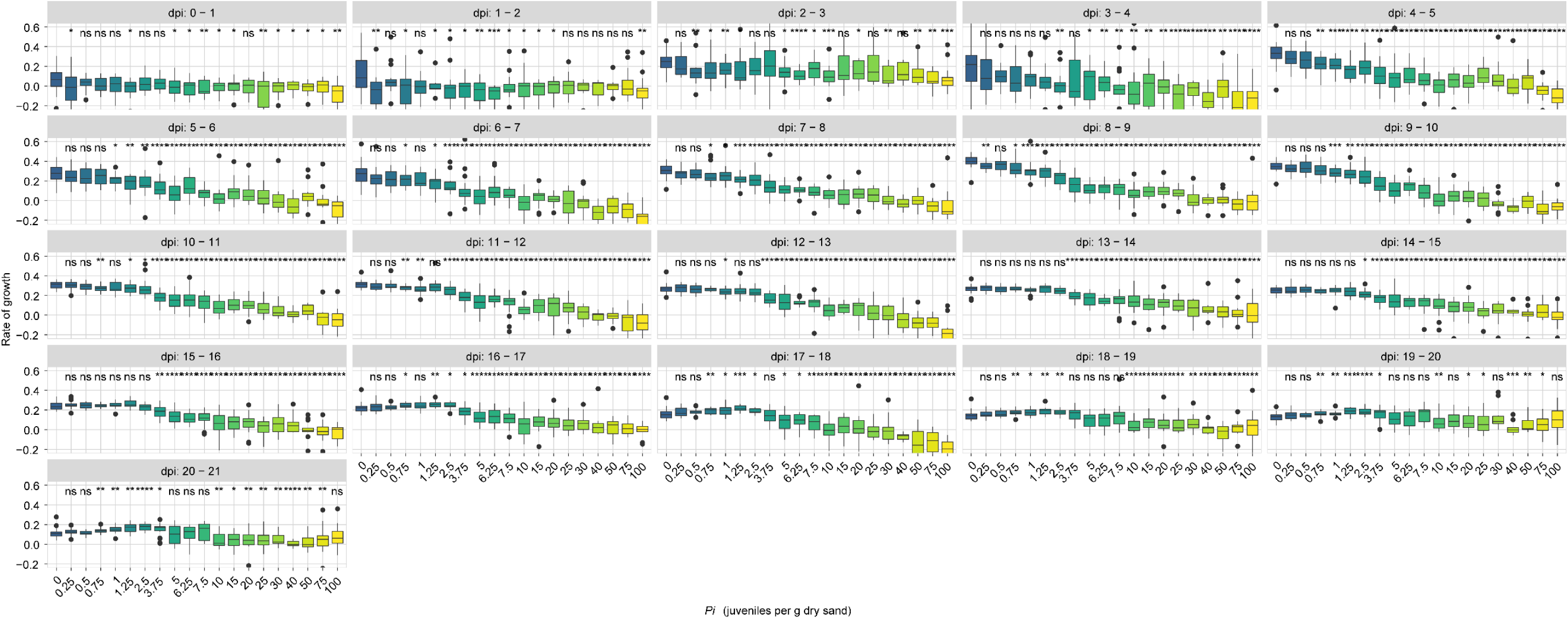
Growth rate of Arabidopsis Col-0 plants inoculated with increasing densities of *Heterodera schachtii*. Nine-day-old Arabidopsis seedlings were inoculated with 20 densities (*Pi*) of *H. schachtii* juveniles (0 to 100 juveniles per g dry sand). The growth rates of plants were calculated per day. Dots represent individual plants. Data was analysed with a Wilcoxon Rank Sum test. ns= not significant, *p< 0.05, **p< 0.01, ***p<0.001 (n=10-24).

**Figure S7:**
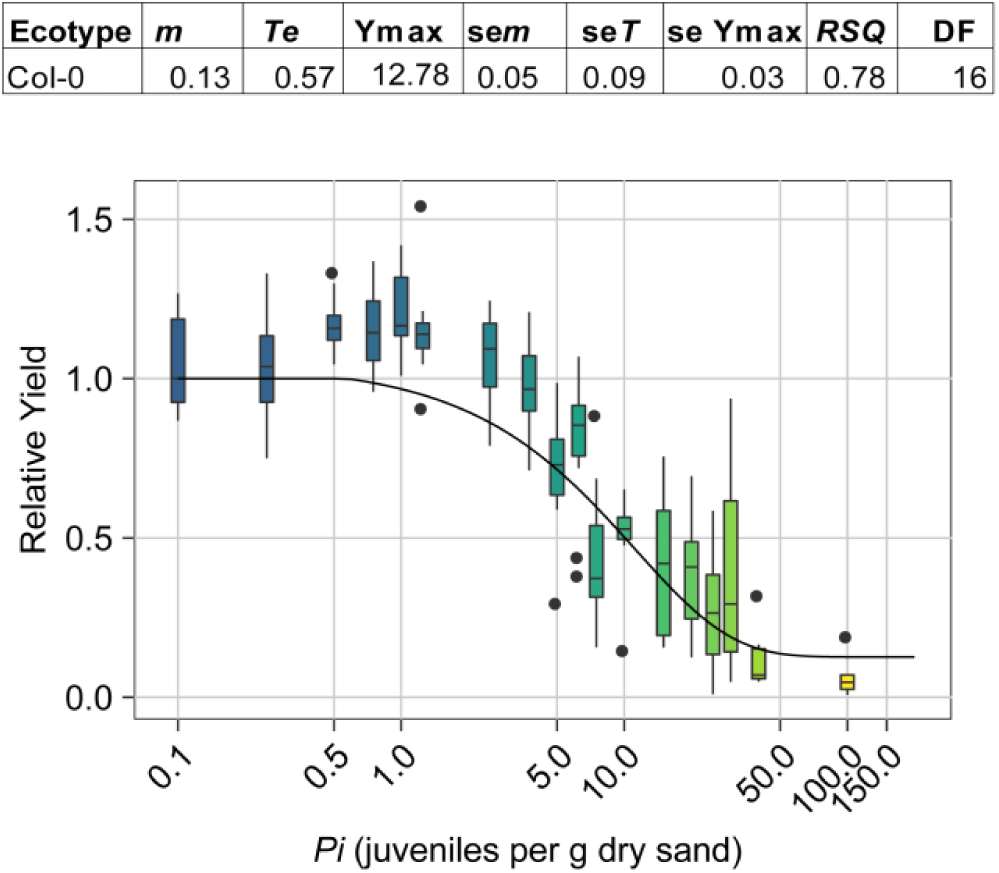
The relationship between the inoculation density (*Pi*) of *Meloidogyne incognita* and green canopy area (Relative Yield). Nine-day-old Arabidopsis seedlings were inoculated with 18 densities of *M. incognita* juveniles (0 to 20.000 J2s per pot) in 200 mL pots containing 200 grams of dry sand. Line was fitted according to the Seinhorst yield loss equation: *y* = *m* + (1 - *m*) 0.95 ^*Pi/T*-1^ for *Pi* > *T* and *y* = 1 for *Pi* ≤ *T*. Parameter values for Seinhorst’s Eq. for the relation between initial population density (*Pi*) of *H. schachtii* and measured leaf surface area. *Pi* and tolerance limit (*T*) are expressed in *M. incognita* (g dry sand)^-1^ while, the minimal yield (*m*) is the lowest proportion of the maximum green canopy area (cm^2^) (*Ymax*) at 21dpi. The goodness of the fit is expressed in the *RSQ*.

**Figure S8:**
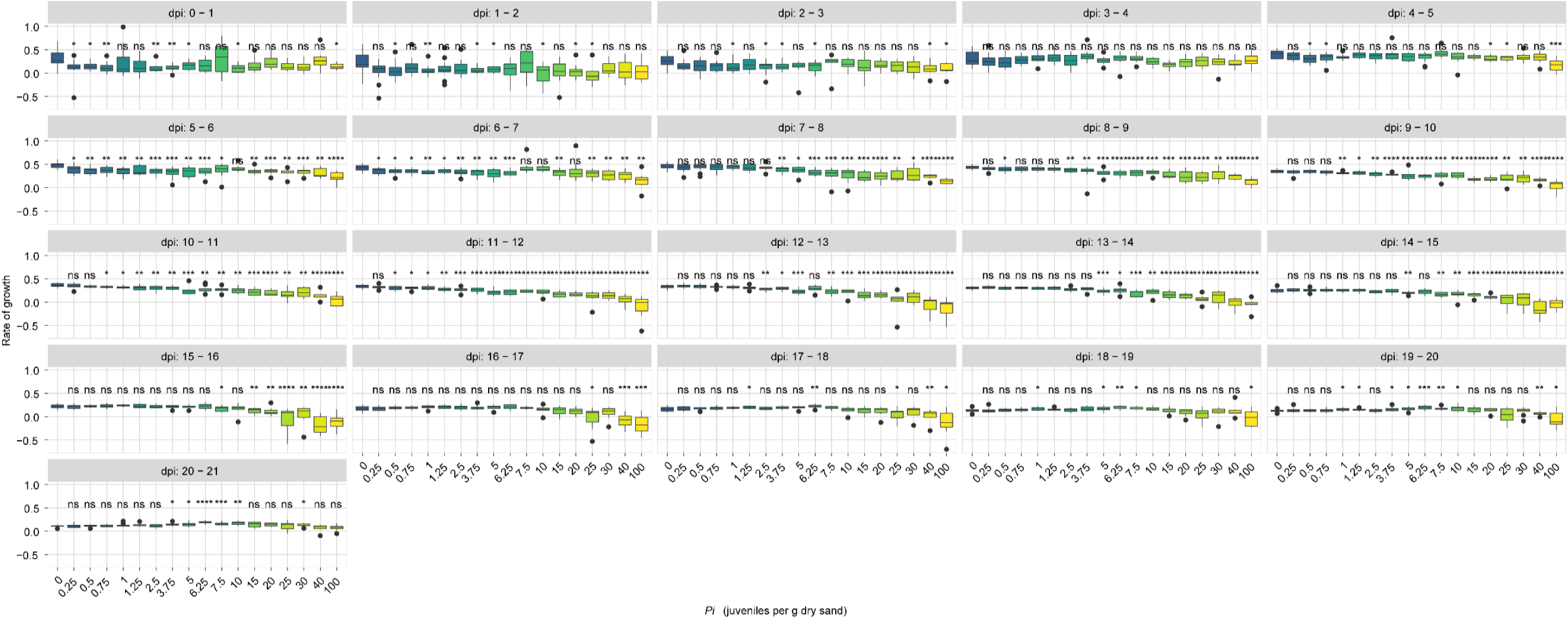
Growth rate of Arabidopsis Col-0 plants inoculated with increasing densities of *Meloidogyne incognita*. Nine-day-old Arabidopsis seedlings were inoculated with 18 densities (*Pi*) of *M. incognita* juveniles (0 to 100 juveniles per g dry sand). The growth rates of plants were calculated per day. Dots represent individual plants. Data was analysed with a Wilcoxon Rank Sum test. ns= not significant, *p< 0.05, **p< 0.01, ***p<0.001 (n=10-12).

**Figure S9:**
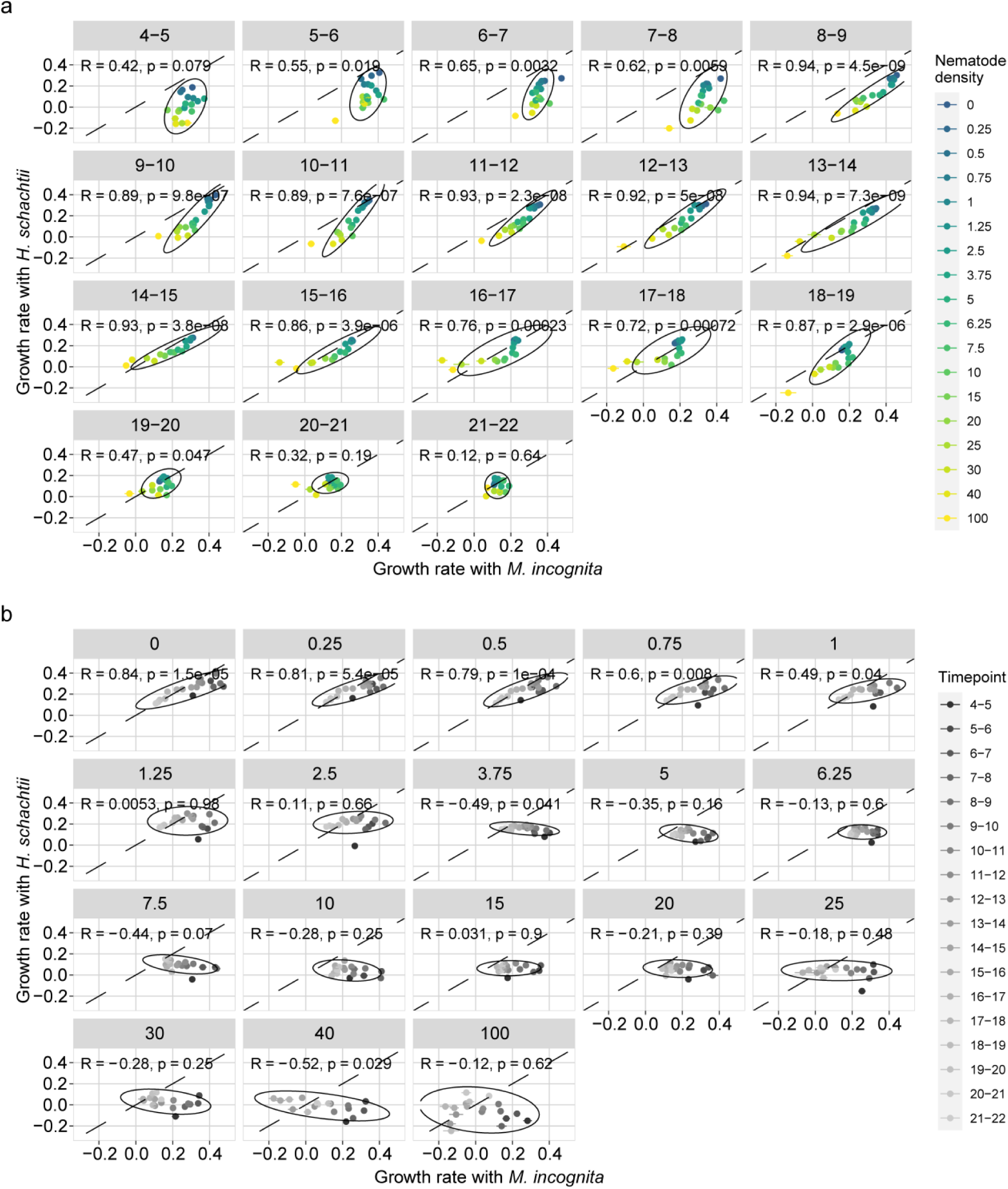
Correlations between growth rates of Arabidopsis Col-0 plants inoculated with *Heterodera schachtii* and *Meloidogyne incognita* per (a) dpi or (b) inoculum density. Nine-day-old Arabidopsis seedlings were inoculated with 18 densities (*Pi*) of *H. schachtii* and *M. incognita* juveniles (0 to 100 juveniles per g dry sand). The growth rates of plants were calculated per day. Dots represent the average growth rate. The correlation coefficient and p-values shown derive from a Pearson correlation.

**Figure S10:**
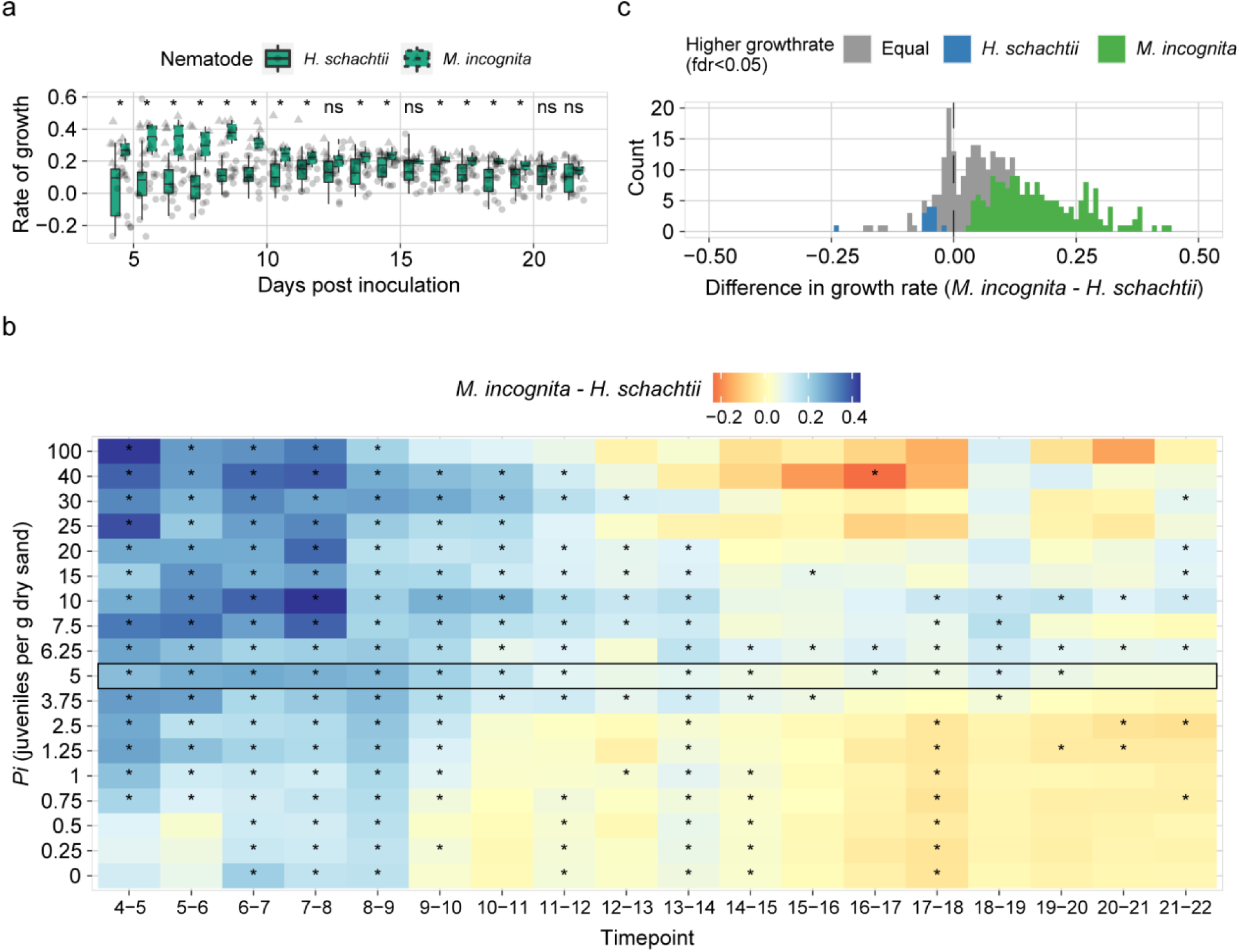
Comparison of growth rates of Col-0 inoculated with *H. schachtii* and *M. incognita*. Arabidopsis seedlings were inoculated with 18 densities (*Pi*) of *H. schachtii* juveniles or 18 *P*_*i*_’s of *M. incognita* (0 to 100 juveniles per g dry sand). Growth rates of green canopy areas were calculated from 0 to 22 dpi. **a)** Comparison of green canopy area growth rates of plants inoculated with *Pi* 5 of *H. schachtii* (full-line boxplots) and *M. incognita* (dashed-line boxplots) over time. **b)** Heatmap of the differential growth rate per *Pi* and timepoint. The difference in growth rate was calculated by subtracting the *H. schachtii* treated growth rates from the *M. incognita* treated growth rates per timepoint and *Pi*, blue indicates that *M. incognita* inoculated plants have a higher growth rate than *H. schachtii* inoculated plants and red indicates vice versa. **c)** Histogram of the number of occurrences that growth rates of *H. schachtii* and *M. incognita* inoculated plants significantly differ per *Pi* and timepoint. Blue represents significantly higher growth rates for *H. schachtii* inoculated plants, grey indicates no significant differences, and green represents significantly higher growth rates for *M. incognita* inoculated plants. Differences in growth rate between plants infected with either *H. schachtii* or *M. incognita* were tested using a paired t-test comparing data from the same day and density combination; ns= not significant, *p< 0.05 (n=10-24).

